# Mitral regurgitation induces a unique fibroblast population associated with atrial fibrillation susceptibility

**DOI:** 10.64898/2026.08.19.745870

**Authors:** Samantha J. Procasky, Jack J. Yi, Emma F. Jones, M. Cassandra Witt, Victoria E. Davis, Alexander N. Wein, Matthew R. Schill, Stacey L. Rentschler, Andrew E. Gelman, Ralph J. Damiano, Christian W. Zemlin

**Author notes:** Corresponding Author: Christian W. Zemlin PhD, Washington University School of Medicine 660 South Euclid Avenue, St. Louis, MO 63110 USA.

## Abstract

**Background:** Mitral regurgitation (MR) is a major risk factor for the development of atrial fibrillation (AF), yet the molecular mechanisms linking volume overload to arrhythmogenic remodeling remain poorly understood. Although fibrosis has long been considered the primary substrate for AF, increasing evidence suggests that fibroblast heterogeneity and cell-cell interactions may play important roles in disease progression.

**Methods:** MR was created endovascularly by chordal avulsion in 12 dogs with 6 controls. AF inducibility was assessed by transvenous burst pacing, left atrial volume by echocardiography, and collagen content by Masson trichrome and picrosirius red staining. Single-nucleus RNA sequencing (snRNA-seq) was performed on left atrial posterior wall tissue from control, 4-week, and 6-month MR animals. Fibroblast subpopulations and fibroblast-cardiomyocyte communication were analyzed and markers validated by RNA in situ hybridization in all 18 animals.

**Results:** MR resulted in progressive left atrial dilation, but neither the change in left atrial volume from baseline nor total collagen burden correlated with the inducibility of AF (n=6 each). SnRNA-seq resolved seven major cardiac cell populations and identified four transcriptionally distinct fibroblast populations (NOX4/GRIA4, PCOLCE2, ADRB2/HCN1, PTX3/ICAM1). Fibroblast composition shifted markedly: matrix-associated PCOLCE2 fibroblasts starkly declined by 6 months, whereas inflammatory-associated PTX3/ICAM1 fibroblasts expanded stepwise over time. Cardiomyocyte-to-fibroblast signaling, dominated by PTPRM and LAMA2, was progressively redirected toward PTX3/ICAM1 fibroblasts. RNAscope confirmed a stepwise rise in ICAM1 transcripts and higher ICAM1 in AF-inducible than non-inducible animals.

**Conclusions:** In a canine model of MR, the inducibility of AF was associated with fibroblast state remodeling rather than with atrial dilation or collagen burden. Progressive expansion of inflammatory-associated PTX3/ICAM1 fibroblasts, together with reorganized fibroblast-cardiomyocyte signaling, defines a candidate arrhythmogenic mechanism and therapeutic target in MR.

## Introduction

Mitral regurgitation (MR), the most common left-sided valvular heart disease in the US, affects up to 2.5 million patients a year in the United States^1^. Atrial fibrillation (AF) is present in up to half of patients with mitral valvular disease^2^, and is independently associated with reduced long-term survival.^3^ Clinical evidence has suggested that treating AF at the time of mitral valve surgery improves survival outcomes^4^. The optimal timing of intervention to prevent irreversible remodeling and AF remains undefined because the underlying mechanisms are poorly understood^5^.

MR produces sustained volume overload and progressive left atrial stretch, initiating structural and molecular remodeling of the atrial myocardium^6,7^. Mechanical stress has been shown in-vitro to directly activate cardiac fibroblasts and stimulate profibrotic signaling pathways^8,9^, leading to extracellular matrix (ECM) remodeling and alterations in tissue architecture. Fibrosis has long been considered a principal substrate for lone AF^10^. However, increasing evidence suggests that fibrosis burden alone does not consistently predict arrhythmia susceptibility^11,12^, but rather the spatial organization and distribution of fibrotic remodeling may have a greater influence on conduction abnormalities^13^.

Cardiac fibroblasts are increasingly recognized as active regulators of myocardial electrophysiology, forming heterocellular gap junctions with cardiomyocytes through connexins, enabling electrical coupling that influences conduction velocity, excitability, and impulse propagation^14–16^. Fibroblasts also modify connexin expression and gap junction organization following cardiac injury^15,17^. Activated and inflammatory fibroblasts located adjacent to cardiomyocytes disrupt connexin localization, impair intercellular coupling, and promote conduction block and reentrant arrhythmias^16,18^. Together, these observations suggest that fibroblasts actively participate in atrial electrical remodeling.

Fibroblast activation, demonstrated through fibroblast expansion, ECM deposition, and electrophysiologic remodeling, has been observed in both experimental and clinical studies with myocardial infarction and pressure overload^19–21^. In contrast, atrial remodeling during MR is less well characterized and may be fundamentally different. Volume overload has been associated with reduced collagen deposition^22^ and increased matrix degradation^23^, but the contribution of fibroblast phenotypic remodeling to arrhythmogenesis remains poorly understood. Consistent with these observations, canine models of MR have exhibited substantial atrial dilation despite relatively modest fibrotic remodeling^6,7,11,12,22,24^, underscoring that AF development in MR likely involves mechanisms beyond collagen accumulation.

Single-nucleus RNA sequencing (snRNA-seq) identifies distinct fibroblast populations and infers intracardiac cell-cell communication while preserving cellular heterogeneity. Using our previously established canine MR model^7,25^, we performed snRNA-seq of the left atrial posterior wall (LAPW) from control, 4-week MR, and 6-month MR animals to characterize cellular remodeling during disease progression. We combined snRNA profiling with in situ RNA hybridization to define fibroblast heterogeneity and its relationship to atrial remodeling. Four distinct fibroblast populations underwent dynamic temporal and regional remodeling throughout MR progression. Importantly, remodeling of the fibroblast populations was associated with inducibility of AF, supporting a multicellular model of arrhythmogenic remodeling involving fibroblast state transitions and altered fibroblast-cardiomyocyte communication.

## Methods

### Animal Choice and MR Surgery

Adult purpose-bred mongrel hounds (25.5 ± 5.7 kg, n=18; 9 male, 9 female) were included in this study. All protocols were reviewed and approved by Washington University in St. Louis IACUC and complied with the NIH Guidelines for Care and Use of Lab Animals. MR was created endovascularly by chordal avulsion under transesophageal echocardiography-guidance^25^. AF inducibility was assessed at baseline and terminal procedures using a transvenous pacing catheter positioned in the right atrial appendage. Among the 12 MR canines, six were classified as AF-positive: five developed sustained AF lasting >300 seconds, and one exhibited spontaneous AF on Holter monitoring (**Figure S1**). The remaining six canines experienced either no induced AF or episodes lasting ≤30 seconds and were classified as AF-negative. Control dogs did not have MR but underwent the same terminal procedure (**Figure 1A**). Six canines each served as controls, 4-week MR, and 6-month MR groups.

**Figure 1.**
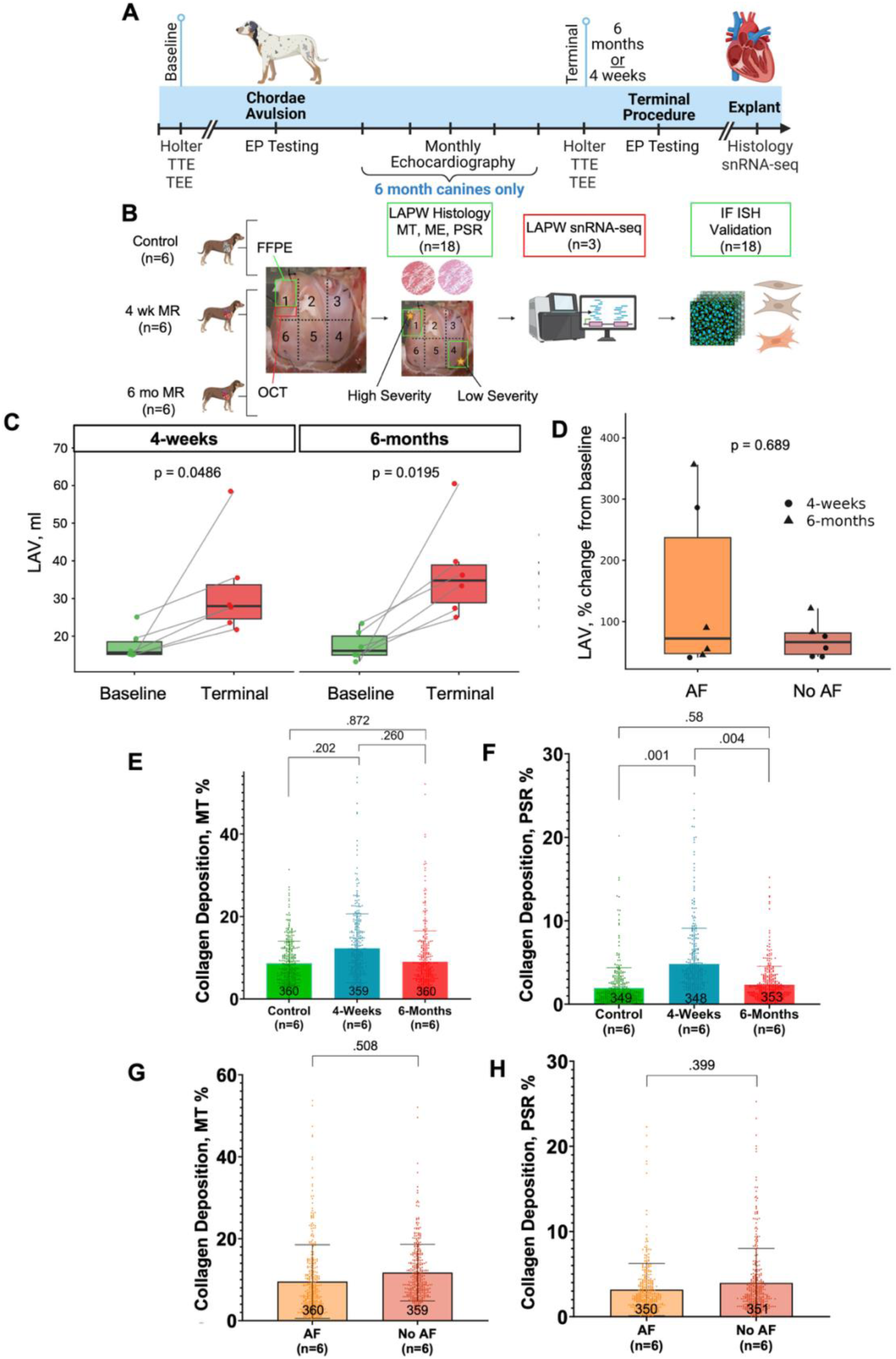
Experimental design and structural remodeling in the canine MR model. **A**, Schematic illustrating induction of mitral regurgitation (MR) and study design. **B**, Histology-guided tissue selection strategy. LAPW regions exhibiting high-and low-remodeling severity were identified and selected for snRNA-seq. **C**, LAV demonstrates progressive atrial enlargement at both 4 weeks MR and 6 months MR (n=6 per group). **D**, Percent change in LAV from baseline stratified by AF inducibility status. Individual datapoints represent animals from the 4-week and 6-month MR cohorts (n=6 per group). **E,** Quantification of collagen deposition from MT-stained sections expressed as the percentage of blue-stained pixels relative to total ROI area. **F,** Quantification of collagen deposition from PSR-stained sections expressed as the percentage of red-stained pixels relative to total ROI area. **G,** Quantification of collagen deposition from MT-stained sections expressed as the percentage of blue-stained pixels relative to total ROI area in animals with No AF and AF. **H,** Quantification of collagen deposition from PSR-stained sections expressed as the percentage of red-stained pixels relative to total ROI area in No AF and AF groups. Values are presented as mean ± SD. Statistical comparisons were performed using linear mixed-effects models.

### Sample Collection and Histology

During the terminal surgery, the heart was arrested with cardioplegia and excised. The LAPW was dissected free and divided into 6 distinct regions (**Figure 1B**). Six samples were selected for snRNA-seq based on AF inducibility and remodeling severity determined by a blinded pathologist using hematoxylin and eosin (HE) and Masson’s trichrome (MT) slides (**Table S1**).

### Collagen Deposition Quantification

Ten randomly chosen transmural regions of interest (ROIs) were selected from each LAPW section from each canine (**Figure S2)**. MT and picrosirius red (PSR) images were analyzed at the ROI level using MATLAB-based image analysis pipelines, as previously described^26^.

### SnRNA-seq

Summary QC metrics are provided in **Figure S3A-F**. The number of nuclei retained for each sample following quality control filtering is provided in **Table S2**. Cell types were annotated using canonical lineage markers identified through differential expression analysis and visualization of cluster-specific marker expression^27^. Major cardiac cell populations were assigned based on established lineage-specific marker expression profiles from CellxGene^28^. Fibroblast subpopulations were subsequently analyzed separately to identify transcriptionally distinct cellular states^29^.

### RNAScope

Spatial validation of candidate fibroblast marker genes identified by snRNA-seq was performed using RNAscope Multiplex Fluorescent In Situ Hybridization (Advanced Cell Diagnostics, Newark, CA, USA) on FFPE left atrial tissue sections^30^. Canine-specific RNAscope probes targeting intercellular adhesion molecule 1 (ICAM1), glutamate ionotropic receptor AMPA type subunit 4 (GRIA4), procollagen C-endopeptidase enhancer 2 (PCOLCE2), and adrenoceptor beta 2 (ADRB2) were used to validate fibroblast subpopulations identified by snRNA-seq.

## Statistical Analysis

All statistical analyses were performed in R version 4.4.0. Statistical tests were selected according to the experimental modality and unit of analysis. Detailed methods are provided in supplemental materials.

## Data Availability

The raw data generated in this study have been deposited in the Zenodo database and the accession code will be added here upon acceptance of the manuscript.

## Code Availability

The ScaleRna Nextflow pipeline for processing raw Fastq reads to feature-barcode matrices are available at Github (https://github.com/ScaleBio/ScaleRna). The code and processed data needed to reproduce the key findings of this paper is found at GitHub (https://github.com/samanthaprocasky/Fibroblasts-MR-2026)

## Results

### Atrial dilation and collagen deposition do not distinguish AF susceptibility

MR resulted in significant LA dilation at both 4 weeks (n=6) and 6 months (n=6) compared to baseline (**Figure 1C**). In the 4-week canines, LAV increased from 15.6 (15.2, 18.5) mL to 28.0 (24.6, 33.7) mL. In the 6-month canines, LAV increased from 16.1 (15.0, 20.0) mL to 34.8 (28.9, 38.9) mL. AF was inducible in 2 of 6 (33%) 4-week and 4 of 6 (67%) 6-month MR canines. Despite the progressive atrial dilation, changes in left atrial volume (LAV) from baseline did not differ between AF inducible (n=6) and non-inducible (n=6) canines (72.4% [47.8%, 237.0%]; 66.5% [46.6%, 81.4%]) (**Figure 1D**).

MT staining showed no significant differences in percent collagen deposition between controls (8.6 ± 5.4%), 4-week MR (12.3 ± 8.4%), and 6-month MR (9.0 ± 7.5%) (**Figure 1E**). PSR staining demonstrated increased percent collagen deposition from controls (1.9 ± 2.4%) to 4-week MR (4.8 ± 4.3%), with no significant difference between controls and 6-month MR (2.3 ± 2.2%). Percent collagen deposition by PSR staining decreased from 4 weeks to 6 months (**Figure 1F**).

However, there were no significant differences in either MT or PSR deposition between the non-inducible (11.7 ± 6.9%; 4.0 ± 4.0%) and AF inducible groups (9.5 ± 9.0%; 3.2 ± 3.1%) (**Figure 1G, H**).

### Fibroblast differentiation is a unique feature of MR

Unique molecular identifiers (UMI) detected per nucleus did not differ between samples, conditions, or severity groups (**Figure S3A–F**). Integrated analysis of the merged (n=6) dataset identified seven major cell types, including adipocytes, macrophages, cardiac neurons, vascular smooth muscle cells, cardiomyocytes, fibroblasts, and endothelial cells, with annotations confirmed by canonical marker expression^27,28^ and distinct cluster-enriched genes. (**Figure 2A-B, Figure S3I-O**). A heatmap of top marker expression demonstrated distinct expression patterns between clusters (**Figure 2C**). There were no significant differences in major cell cluster proportions between conditions or severity groups (**Figure 2D, E**). Within the fibroblast cluster, cell composition differed between conditions (**Figure 2F**), samples, and severity groups (**Figure S3G, H**), which led us to investigate this population further.

**Figure 2.**
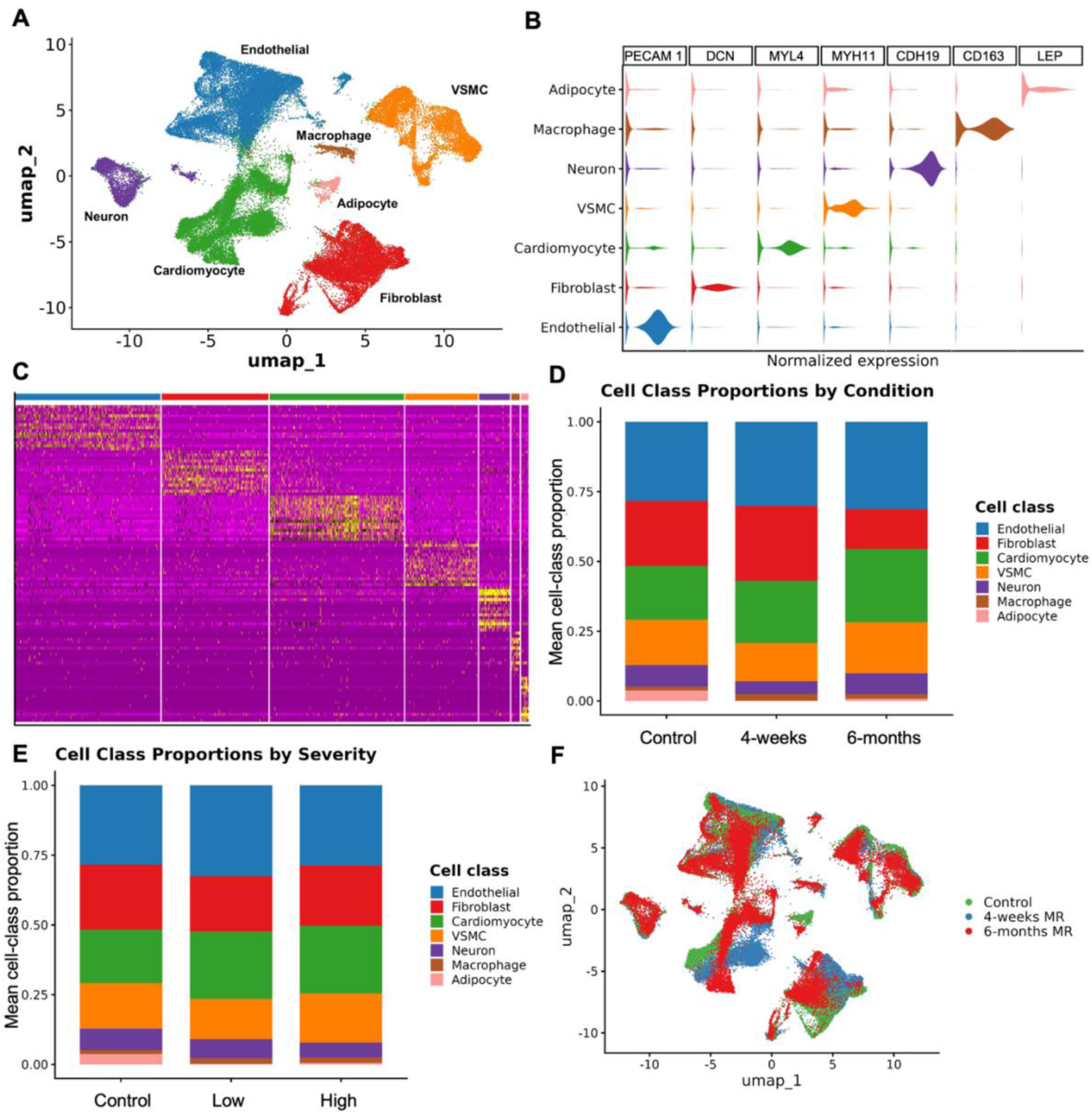
Integrated snRNA-seq characterization of LA remodeling in MR. **A**, UMAP visualization from six LAPW samples. **B**, Expression of canonical marker genes used for cell-type annotation. **C**, Heatmap of the top differentially expressed marker genes defining each major cell population. **D**, Sample-level proportions of major cardiac cell populations across conditions. **E**, Sample-level proportions across remodeling severity groups. Each point represents one biological sample, and values are presented as median (interquartile range [IQR]). Overall differences were assessed using Kruskal-Wallis tests followed by pairwise Wilcoxon rank-sum tests. **F**, UMAP visualization colored by experimental condition demonstrating shifts in cellular distributions.

We identified four LAPW fibroblast subsets, distinguished by differential transcript accumulation of NOX4/GRIA4, PCOLCE2, ADRB2/HCN1, or PTX3/ICAM1 (**Figures 3A & B**). Notably, these subsets did not exhibit potential differences in proliferation, as cell-cycle phase-associated transcript levels were similar across clusters (**Figure S4A**). However, each fibroblast population displayed distinct marker expression profiles, shown by heatmap and feature plots (**Figure S4B–J**). NOX4/GRIA4 fibroblasts showed enrichment of transcripts that promote cell adhesion, extracellular matrix/structural organization, and collagen fibril organization (**Figure 3C**), while PCOLCE2 fibroblasts exhibited potential for enzyme-linked receptor protein and transmembrane receptor protein tyrosine kinase signaling pathways (**Figure 3D**). In contrast, ADRB2/HCN1 fibroblasts revealed enrichment of transcripts that encode for regulation of protein-containing complex assembly, regulation of canonical Wnt signaling, and negative regulation of Wnt signaling pathways (**Figure 3E**). Wnt signaling is dysregulated in atrial tissue from patients with AF, including non-canonical Wnt signaling components^31^. In cardiac fibroblasts more broadly, Wnt signaling regulates fibroblast migration and myofibroblast differentiation, participates in cardiac fibrosis, and promotes fibroblast inflammatory responses in heart failure^32,33^. Expression of ADRB2 and HCN1 further suggests responsiveness to β-adrenergic signaling and ion channel-associated regulatory pathways that may influence fibroblast-cardiomyocyte communication. PTX3/ICAM1 fibroblasts showed enrichment of amide metabolic processes, peptide metabolic processes, and positive regulation of miRNA metabolic process pathways (**Figure 3F**).

**Figure 3.**
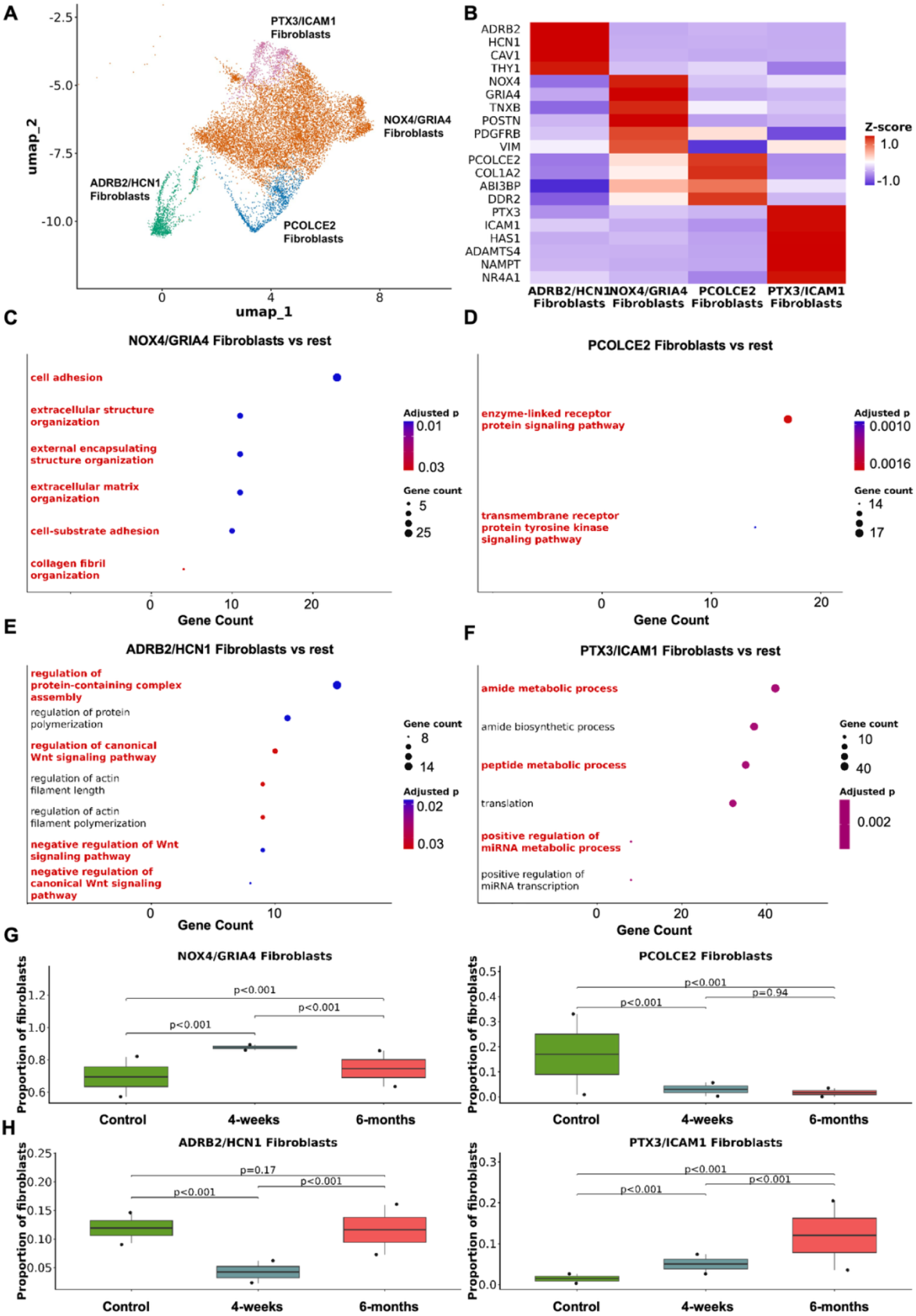
Fibroblast cell state diversification in the volume-overloaded heart **A**, UMAP visualization of fibroblasts isolated from the LAPW revealing four transcriptionally distinct fibroblast populations: NOX4/GRIA4, PCOLCE2, ADRB2/HCN1, and PTX3/ICAM1 fibroblasts. **B**, Heatmap showing scaled expression of canonical marker genes used to define each fibroblast population. **C-F**, GO enrichment analysis of genes upregulated in each fibroblast population compared with remaining fibroblasts. Dot size corresponds to the number of genes associated with each pathway and color represents the p-value. **G, H,** Sample-level fibroblast subpopulation proportions across experimental conditions. Each point represents one biological sample. Statistical comparisons were performed using Kruskal-Wallis tests followed by pairwise Wilcoxon rank-sum tests.

Fibroblast subpopulation proportions differed significantly across conditions (**Figure 3G, H; Table 1**). NOX4/GRIA4 fibroblasts increased at 4 weeks and 6 months compared with controls and declined from 4 weeks to 6 months. PCOLCE2 fibroblasts decreased at both 4 weeks and 6 months compared with controls, with no difference between the two MR time points. ADRB2/HCN1 fibroblasts decreased at 4 weeks compared with controls, returned to control levels by 6 months, and increased between 4 weeks and 6 months. In contrast, PTX3/ICAM1 fibroblasts showed a progressive increase throughout disease progression, increased at both 4 weeks and 6 months of MR compared to controls and further increased from 4 weeks to 6 months in the MR cohort.

**Table 1.**
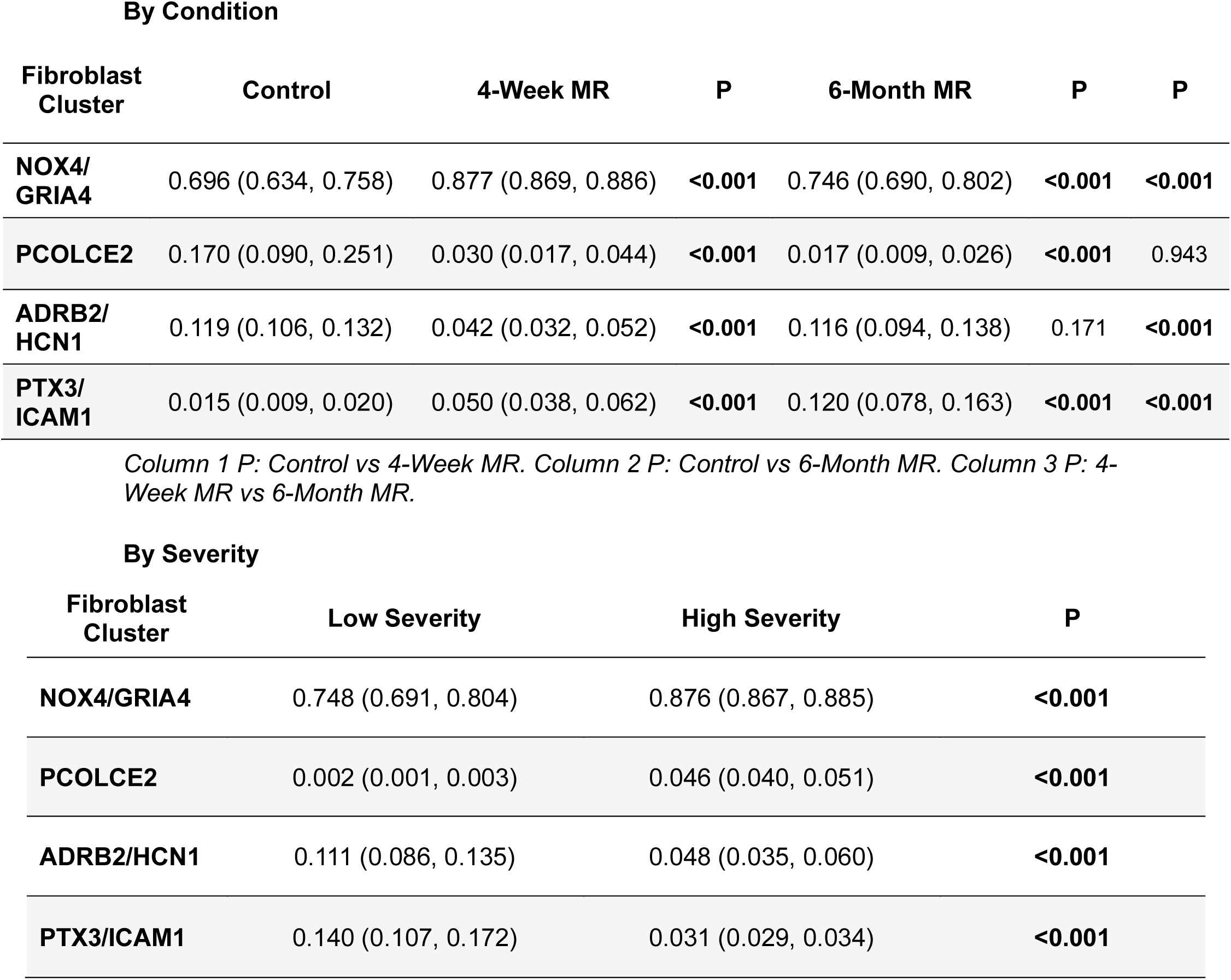
snRNA-seq fibroblast cluster proportions by condition and severity. Values are presented as median (IQR) fibroblast cluster proportions within the total fibroblast population for each sample, as determined by snRNA-seq. P values were calculated using Wilcoxon rank-sum tests.

**By Condition**
| Fibroblast Cluster | Control | 4-Week MR | P | 6-Month MR | P | P |
| --- | --- | --- | --- | --- | --- | --- |
| NOX4/GRIA4 | 0.696 (0.634, 0.758) | 0.877 (0.869, 0.886) | <0.001 | 0.746 (0.690, 0.802) | <0.001 | <0.001 |
| PCOLCE2 | 0.170 (0.090, 0.251) | 0.030 (0.017, 0.044) | <0.001 | 0.017 (0.009, 0.026) | <0.001 | 0.943 |
| ADRB2/HCN1 | 0.119 (0.106, 0.132) | 0.042 (0.032, 0.052) | <0.001 | 0.116 (0.094, 0.138) | 0.171 | <0.001 |
| PTX3/ICAM1 | 0.015 (0.009, 0.020) | 0.050 (0.038, 0.062) | <0.001 | 0.120 (0.078, 0.163) | <0.001 | <0.001 |

| Fibroblast Cluster | Low Severity | High Severity | P |
| --- | --- | --- | --- |
| NOX4/GRIA4 | 0.748 (0.691, 0.804) | 0.876 (0.867, 0.885) | <0.001 |
| PCOLCE2 | 0.002 (0.001, 0.003) | 0.046 (0.040, 0.051) | <0.001 |
| ADRB2/HCN1 | 0.111 (0.086, 0.135) | 0.048 (0.035, 0.060) | <0.001 |
| PTX3/ICAM1 | 0.140 (0.107, 0.172) | 0.031 (0.029, 0.034) | <0.001 |

**Table 2.** RNAscope validation of fibroblast population marker expression across condition, AF status, and severity. Values are presented as median (IQR) for total RNAscope puncta counts per ROI. Pairwise comparisons were performed using Wilcoxon rank-sum tests. Data are shown for the four fibroblast population markers selected for validation from the snRNA-seq analysis: PCOLCE2, GRIA4, ICAM1, and ADRB2.

**By Condition**
| Marker | Control (n=6) | 4-week MR (n=6) | P | 6-month MR (n=6) | P | P |
| --- | --- | --- | --- | --- | --- | --- |
| PCOLCE2 | 42.00 (12.50, 81.50) | 14.00 (7.25, 40.50) | .010 | 18.50 (11.00, 40.75) | .020 | .286 |
| GRIA4 | 0.00 (0.00, 0.00) | 0.00 (0.00, 1.00) | .033 | 0.00 (0.00, 0.00) | .111 | .662 |
| ICAM1 | 69.00 (47.50, 103.50) | 116.00 (58.75, 329.50) | <.001 | 312.50 (234.00, 768.50) | <.001 | <.001 |
| ADRB2 | 23.00 (6.50, 40.00) | 28.50 (17.00, 42.25) | .098 | 49.00 (31.00, 66.50) | <.001 | <.001 |

**By AF Status**
| Marker | AF | No AF | P |
| --- | --- | --- | --- |
| PCOLCE2 | 28.00 (11.00, 45.00) | 14.00 (6.00, 25.50) | .015 |
| GRIA4 | 0.00 (0.00, 0.00) | 0.00 (0.00, 1.00) | .370 |
| ICAM1 | 328.00 (252.00, 831.00) | 109.00 (59.50, 312.50) | <.001 |
| ADRB2 | 48.00 (28.00, 67.00) | 30.00 (18.50, 45.00) | .002 |

**By Severity**
| Marker | Control | Low Severity | P | High Severity | P | P |
| --- | --- | --- | --- | --- | --- | --- |
| PCOLCE2 | 42.00 (12.50, 81.50) | 14.00 (8.00, 30.00) | <.001 | 24.00 (11.50, 44.50) | .034 | .059 |
| GRIA4 | 0.00 (0.00, 0.00) | 0.00 (0.00, 0.00) | .685 | 0.00 (0.00, 2.50) | .002 | .006 |
| ICAM1 | 69.00 (47.50, 103.50) | 277.00 (138.00, 405.00) | <.001 | 274.00 (105.50, 426.50) | <.001 | .774 |
| ADRB2 | 23.00 (6.50, 40.00) | 31.00 (23.00, 58.00) | .002 | 39.00 (23.00, 59.50) | <.001 | .362 |

Column 1 P: Control vs Low Severity. Column 2 P: Control vs High Severity. Column 3 P: Low Severity vs High Severity.

Fibroblast subpopulation proportions also differed significantly between low-and high-severity remodeling regions (**Figure S4I; Table 1**). NOX4/GRIA4 and PCOLCE2 fibroblasts were enriched in high-remodeling regions compared with low-remodeling regions. In contrast, ADRB2/HCN1 and PTX3/ICAM1 fibroblasts were reduced in high-severity regions. Based on their cluster-specific expression and minimal expression across other major cell types, ICAM1, GRIA4, ADRB2, and PCOLCE2 were selected for RNAscope validation (**Figure S4K**).

### Inferred fibroblast differentiation patterns and communication networks reveal potential bidirectional communication with cardiomyocytes

Trajectory and pseudotime analysis showed progression of fibroblast states from PCOLCE2 to NOX4/GRIA4, followed by PTX3/ICAM1 and ADRB2/HCN1 fibroblasts (**Figure 4A, B**). Gene ontology (GO) pathway analysis of total fibroblasts comparing 6-month MR with control showed upregulation of intracellular signaling cassette, circulatory system development, vasculature development, blood vessel development, blood vessel morphogenesis, and angiogenesis pathways (**Figure 4C**).

**Figure 4.**
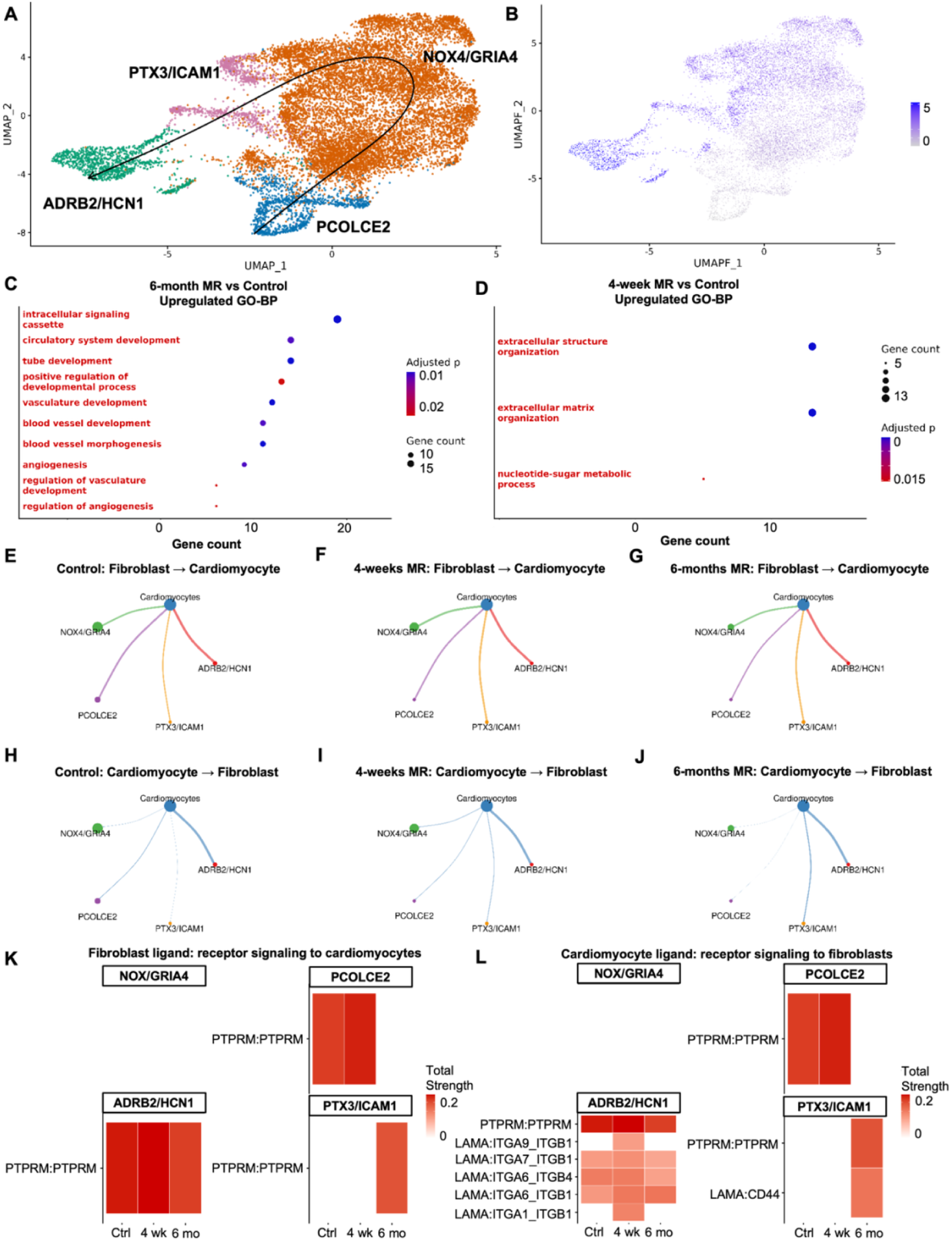
Fibroblast state transitions and fibroblast-cardiomyocyte communication during MR progression. **A**, Trajectory inference analysis of fibroblast populations with arrows indicating the predicted direction of cellular state transitions. **B**, Pseudotime projection illustrating the continuum of fibroblast state progression identified by trajectory analysis. **C, D,** GO analysis of differentially expressed genes in fibroblasts comparing 6-month MR versus control (**C**), and 4-week MR versus control (**D**). Dot size represents the number of genes associated with each pathway, and color indicates adjusted enrichment significance. **E-J**, CellChat network analysis of fibroblast-to-cardiomyocyte signaling interactions in control (**E**), 4-week MR (**F**), and 6-month MR (**G**) samples. Circle plots depict the inferred intercellular communication network, with edge thickness corresponding to interaction strength. **H-J,** CellChat network analysis of cardiomyocyte-to-fibroblast signaling interactions in control (**H**), 4-week MR (**I**), and 6-month MR (**J**) samples. Circle plots illustrate the relative strength and directionality of inferred ligand-receptor signaling between cell populations. **K**, Heatmap of fibroblast-derived ligand-receptor signaling directed toward cardiomyocytes across experimental groups. **L,** Heatmap of cardiomyocyte-derived ligand-receptor signaling directed toward fibroblasts across experimental groups. Heatmap color intensity reflects the relative strength of inferred signaling interactions.

Cell-cell communication analysis revealed differences in fibroblast-to-cardiomyocyte signaling among the control, 4-week, and 6-month conditions (**Figure 4E–G**). In controls, signaling strengths from fibroblasts to cardiomyocytes were 0.92 for NOX4/GRIA4, 1.00 for PCOLCE2, 1.30 for ADRB2/HCN1, and 0.68 for PTX3/ICAM1 fibroblasts (**Figure S5A**). At 4 weeks, signaling strengths were 1.10 for NOX4/GRIA4, 0.90 for PCOLCE2, 1.30 for ADRB2/HCN1, and 0.98 for PTX3/ICAM1 fibroblasts (**Figure S5B**). At 6 months, signaling strengths were 0.52 for NOX4/GRIA4, 0.45 for PCOLCE2, 0.77 for ADRB2/HCN1, and 0.54 for PTX3/ICAM1 fibroblasts (**Figure S5C**).

Cell-cell communication analysis also showed differences in cardiomyocyte-to-fibroblast signaling between conditions (**Figure 4H–J**). In controls, cardiomyocyte-to-fibroblast signaling strengths were 0.084 for NOX4/GRIA4, 0.30 for PCOLCE2, 1.30 for ADRB2/HCN1, and 0.037 for PTX3/ICAM1 fibroblasts (**Figure S5E**). At 4 weeks, signaling strengths were 0.40 for NOX4/GRIA4, 0.35 for PCOLCE2, 2.20 for ADRB2/HCN1, and 0.47 for PTX3/ICAM1 fibroblasts (**Figure S5F**). At 6 months, signaling strengths were 0.012 for NOX4/GRIA4, 0.0083 for PCOLCE2, 1.10 for ADRB2/HCN1, and 0.50 for PTX3/ICAM1 fibroblasts (**Figure S5G**).

Ligand-receptor interactions between fibroblast subpopulations and cardiomyocytes are summarized in **Figure 4K, L,** and **Table S3**. PTPRM-mediated signaling emerged as the predominant fibroblast-cardiomyocyte interaction throughout disease progression, where PTPRM is a contact-dependent receptor protein tyrosine phosphatase that accumulates at sites of cell-cell interaction and regulates intracellular signaling through dephosphorylation of adhesion-associated proteins^34–36^. No ligand-receptor interactions with communication strengths >0.1 were detected between NOX4/GRIA4 fibroblasts and cardiomyocytes under any condition. PTPRM signaling was detected in PCOLCE2 and ADRB2/HCN1 fibroblasts in control tissue and at 4 weeks, whereas at 6 months it persisted only in ADRB2/HCN1 fibroblasts and newly emerged in PTX3/ICAM1 fibroblasts.

Reciprocal cardiomyocyte-to-fibroblast signaling demonstrated a similar pattern, with no interactions detected for NOX4/GRIA4 fibroblasts. In addition to conserved PTPRM signaling, LAMA2-mediated interactions were predominantly directed toward ADRB2/HCN1 fibroblasts across disease progression and toward PTX3/ICAM1 fibroblasts during MR remodeling. Laminin-integrin signaling regulates cell adhesion, mechanosignaling, and extracellular matrix organization in the healthy myocardium, whereas CD44 signaling has been linked to inflammatory fibroblast activation and TGF-β-mediated myofibroblast differentiation^37–39^.

### In situ hybridization validates fibroblast markers in a larger cohort of MR and inducible animals

RNAscope provided independent spatial validation of the fibroblast remodeling patterns identified by snRNA-seq in both the sequenced canines and the larger cohort of 18 animals (**Figure 5A**), demonstrating that these transcriptionally defined fibroblast populations represent reproducible biological remodeling programs.

**Figure 5.**
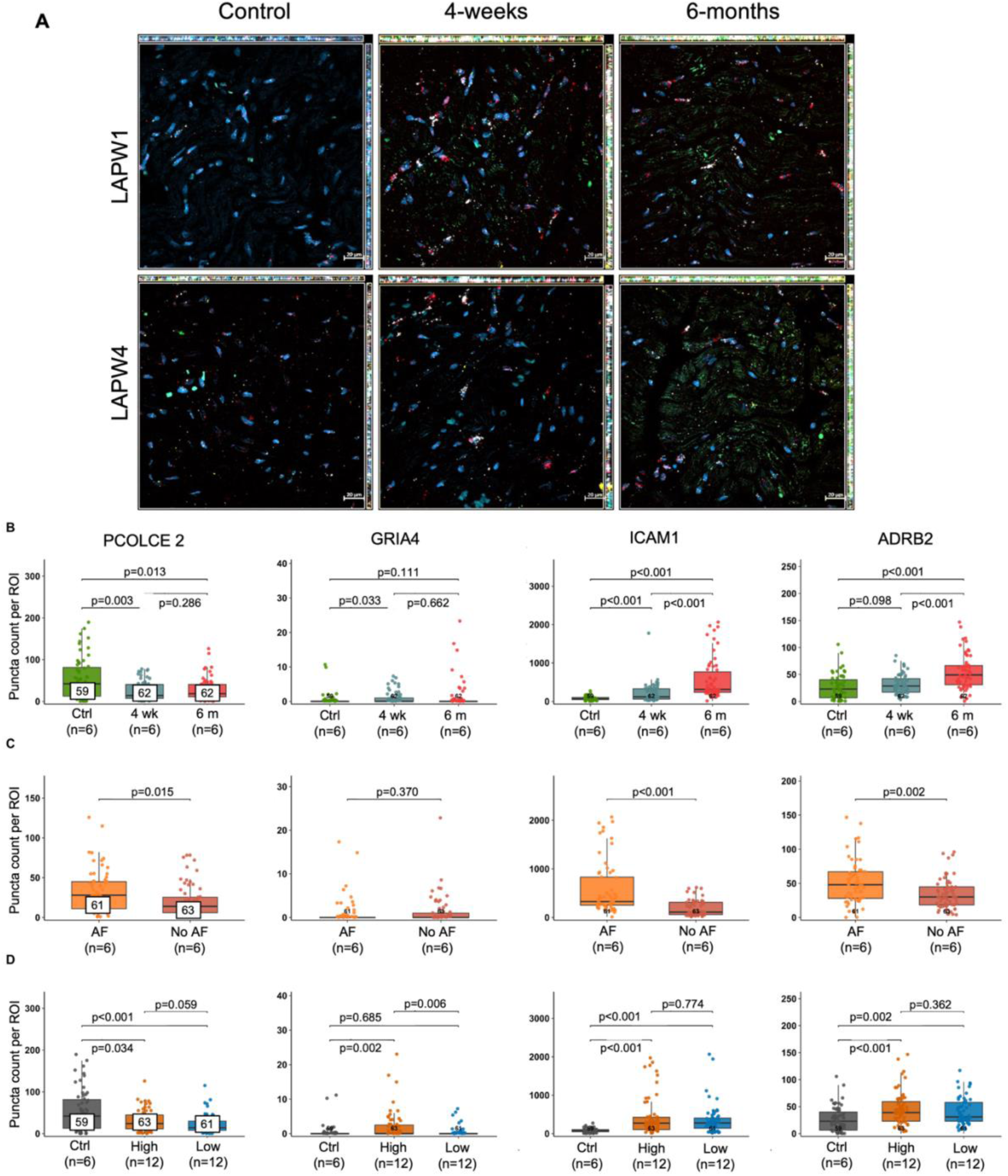
RNAscope validation of fibroblast marker expression across MR progression and AF susceptibility. **A**, Representative 63X RNAscope immunofluorescence images from the six LAPW samples subjected to snRNA-seq. Nuclei are labeled with DAPI (blue). Fibroblast marker transcripts are visualized as ICAM1 (red, Opal 650), GRIA4 (green, Opal 520), ADRB2 (yellow, Opal 570), and PCOLCE2 (white, Opal 780). **B**, Quantification of total RNAscope puncta counts by condition (n=6 each group). ROIs listed in first panel are representative across each row. **C**, Quantification by AF inducibility (n=6 each group). **D**, Quantification by remodeling severity (n=18 total). Individual data represent ROIs, with five ROIs analyzed per tissue section. Pairwise comparisons were performed using Wilcoxon rank-sum tests.

Analysis of the six samples from the three canines used for snRNA-seq demonstrated expression patterns that were consistent with those observed in the larger 18-canine validation cohort (**Figure S6A, B**). Specifically, PCOLCE2 expression decreased at both 4-weeks MR and 6-months MR compared to controls. GRIA4 showed only a modest increase at 4-weeks compared to control. Although the NOX4/GRIA4 population was the largest fibroblast population identified by snRNA-seq, GRIA4 puncta counts remained low by RNAscope. This discrepancy likely reflects the fact that GRIA4 serves as a cluster-defining marker rather than being uniformly expressed across all cells within the transcriptionally defined population. ADRB2 expression increased at 6 months compared to control and 4-week MR. Most notably, ICAM1 demonstrated the most pronounced temporal remodeling, increasing progressively from control to 4-week and 6-month MR, with significant differences between all three groups (**Figure 5B**; **Table 1**).

When stratified by AF inducibility, ICAM1 expression was higher in AF inducible animals than in non-inducible animals (**Figure 5C**; **Table 1**). ADRB2 and PCOLCE2 puncta counts were also increased in AF animals. GRIA4 did not differ between groups. While collagen deposition declined from 4 weeks to 6 months, ICAM1 and ADRB2 expression continued to increase during the same period, coinciding with greater AF inducibility in the MR animals.

Regional analyses demonstrated persistent remodeling across both low-and high-severity tissue regions (**Figure 5D**; **Table 1**). ICAM1 remained significantly elevated in both low-and high-severity regions compared with control tissue, although no difference was observed between severity groups. Similarly, ADRB2 expression was increased in both severity groups, while PCOLCE2 expression was reduced in both low-and high-severity regions relative to controls. GRIA4 expression was increased only in high-severity regions compared with control tissue.

## Discussion

This study provides one of the first comprehensive characterizations of the molecular and cellular remodeling during the progression of MR and its relationship to the development of AF. Our findings suggest that remodeling of distinct fibroblast states may play an important role in the formation of the arrhythmogenic substrate. However, this remodeling response did not result in either increased fibrosis or collagen deposition on histological examination. In fact, although MR produced substantial LA dilation, neither atrial enlargement nor total collagen burden distinguished AF inducible from non-inducible animals. Instead, snRNA-seq identified four transcriptionally distinct fibroblast populations that underwent dynamic temporal and regional remodeling throughout disease progression, with the PTX3/ICAM1 fibroblast population specifically demonstrating strong associations with both duration of MR and AF inducibility. Importantly, these transcriptional programs were independently validated by in situ hybridization in a larger canine cohort. Together, these findings support a model in which specialized fibroblast populations and their potential interactions with cardiomyocytes contribute to atrial remodeling through mechanisms that extend beyond the extent of fibrosis and atrial dilation alone.

MR resulted in significant LA dilation at both 4 weeks and 6 months, consistent with the clinical remodeling in MR^7^. However, the change in LA volume from baseline did not correlate with AF inducibility, indicating that LA dilation does not fully account for arrhythmia vulnerability in this model. While PSR staining showed increased collagen deposition at 4 weeks compared with controls, this increase was not maintained for 6 months. PSR is sensitive to collagen organization and early collagen deposition^40^, which may explain its detection of early matrix remodeling that was not identified by MT. In addition, collagen deposition did not differ between inducible and non-inducible animals. These findings suggest that total collagen burden alone does not explain or correlate with the electrophysiologic remodeling in this MR model and supports further evaluation of non-fibrotic mechanisms, including fibroblast-cardiomyocyte coupling, connexin remodeling, or fibroblast-induced changes in the extracellular matrix.

The abundance of major cardiac cell types remained stable throughout disease progression; however, fibroblast composition changed substantially across MR timepoints and regional severities. These findings suggest that arrhythmogenic remodeling in MR is driven less by large shifts in overall cellular composition than by remodeling within specialized fibroblast populations, highlighting the importance of cell-state transitions rather than broad tissue-type changes. Although multiple transcriptomics studies have demonstrated fibroblast heterogeneity across the healthy and diseased heart, the specific fibroblast populations identified have varied substantially depending on species, cardiac chamber, disease context, and analytical approach. Previous studies have described resident, activated, myofibroblast, and matrifibroblast populations, but none have identified the same marker-defined populations observed in the present study^19,41–43^. Accordingly, we identified four transcriptionally distinct fibroblast populations in the MR LAPW whose molecular programs suggest complementary roles in extracellular matrix maintenance, oxidative stress, inflammation, and cardiomyocyte communication.

The PTX3/ICAM1 population exhibited features of an inflammatory, stress-responsive fibroblast state that may promote atrial remodeling rather than structural fibrosis alone. Given established roles for PTX3 and ICAM1 in inflammatory signaling, leukocyte recruitment, tissue repair, and extracellular matrix remodeling^44–46^, this population may help sustain a pro-inflammatory microenvironment by coordinating immune cell recruitment, amplifying inflammatory signaling, and remodeling the extracellular matrix through paracrine signaling and matrix degradation. Persistent inflammatory fibroblast activation has increasingly been recognized as contributing to atrial electrical remodeling through immune-fibroblast-cardiomyocyte crosstalk, connexin remodeling, conduction heterogeneity, and arrhythmia susceptibility^47^. Experimental studies have further demonstrated that inflammatory signaling is not merely a consequence of AF but can actively promote atrial arrhythmogenesis, with suppression of inflammatory pathways reducing atrial electrical remodeling and AF susceptibility^48–50^. These observations raise the possibility that therapeutic strategies targeting inflammatory fibroblast activation or interrupting fibroblast-mediated inflammatory signaling could mitigate the development of the arrhythmogenic substrate during MR. This population progressively expanded throughout MR progression, demonstrated the strongest RNAscope validation through ICAM1 expression, and was most strongly associated with AF inducibility. Notably, the PTX3/ICAM1 population is consistent with recent work by Simon-Chica et al.^51^, who identified a PTX3-positive fibroblast population enriched within AF driver regions. Together, these findings suggest that inflammatory PTX3-expressing fibroblasts may represent one of the strongest cellular correlates of arrhythmogenic remodeling.

The PCOLCE2 fibroblast population was consistent with a resident fibroblast population involved in collagen deposition and extracellular matrix organization^52,53^. This population declined rapidly following MR induction and remained suppressed throughout disease progression, suggesting an early loss of normal matrix-maintaining fibroblast function, consistent with our histological findings of no increased collagen deposition in MR animals when compared to control at 6 months.

The NOX4/GRIA4 fibroblast population appeared to represent a stress-responsive remodeling state activated by mechanical stretch and oxidative stress, a well-established driver of fibroblast activation, TGF-β signaling, and atrial remodeling^54,55^. Its enrichment for extracellular matrix organization, collagen fibril organization, and cell adhesion pathways suggests that an oxidative stress-associated remodeling program may prime the atrial microenvironment for subsequent inflammatory and electrophysiologic remodeling, representing a transitional state that initiates structural adaptation following injury.

The ADRB2/HCN1 fibroblast population’s transcriptional profile suggested a potential role in electrophysiologic remodeling^56,57^, consistent with evidence that fibroblasts can influence conduction, resting membrane potential, repolarization, excitability, and arrhythmia susceptibility^51,58,59^. This population was associated with AF inducibility, and functional studies are needed to clarify its contribution. Both PTX3/ICAM1 and ADRB2/HCN1 fibroblast populations were reduced within histologically defined high-remodeling severity regions. This may indicate that these fibroblast states preferentially localize to transitional remodeling zones or reflect dynamic regional state transitions as remodeling progresses.

Trajectory and pseudotime analyses suggested a continuum of fibroblast state transitions beginning with the resident PCOLCE2 population, progressing through the stress-responsive NOX4/GRIA4 population, and subsequently transitioning toward the inflammatory PTX3/ICAM1 and remodeling-associated ADRB2/HCN1 populations. Although pseudotime analysis cannot establish lineage relationships^60^, these findings support a model in which MR promotes progressive fibroblast state transitions rather than static expansion of independent fibroblast populations.

Consistent with this model, fibroblasts from the 6-month MR cohort demonstrated enrichment of signaling, vasculature development, angiogenesis, and blood vessel morphogenesis pathways, suggesting that volume overload was accompanied by increasingly complex transcriptional programs beyond ECM remodeling alone.

Although these pathway enrichments suggest complex fibroblast remodeling during MR, additional lineage-tracing and functional studies will be required to establish the biological significance of these predicted state transitions.

Changes in fibroblast state were paralleled by dynamic alterations in predicted fibroblast-cardiomyocyte communication. While fibroblast-to-cardiomyocyte signaling remained relatively stable across disease progression, cardiomyocyte-to-fibroblast communication demonstrated more pronounced temporal remodeling, with early signaling toward NOX4/GRIA4 and ADRB2/HCN1 fibroblasts at 4 weeks and progressively greater communication with PTX3/ICAM1 fibroblasts during disease.

Cardiomyocytes may actively participate in directing fibroblast state transitions during MR progression, resulting in reciprocal remodeling of both cellular compartments rather than fibroblast activation occurring in isolation.

These findings suggest that the remodeling of specific fibroblast populations, particularly inflammation-associated PTX3/ICAM1 fibroblasts and remodeling-associated ADRB2/HCN1 fibroblasts, may contribute to the development of the arrhythmogenic substrate than total extracellular matrix accumulation alone. Rather than supporting fibrosis as the primary determinant of arrhythmia susceptibility, these data support a model in which progressive transitions in fibroblast states and altered fibroblast-cardiomyocyte interactions accompany the development of an arrhythmogenic atrial microenvironment during MR. Future studies should determine whether modulating inflammatory fibroblast activation or interrupting fibroblast-cardiomyocyte signaling can alter fibroblast state transitions and ultimately reduce susceptibility to AF during MR.

## Conclusion

This study provides one of the first comprehensive characterization of cellular and fibroblast heterogeneity in a model of MR and correlated these changes with susceptibility to AF. Although MR produced progressive left atrial dilation, neither atrial enlargement nor total collagen burden alone explained AF inducibility. Instead, we identified dynamic changes in transcriptionally distinct fibroblast populations that occurred throughout disease progression, varied across regional severities, and were validated by RNAscope in an independent cohort.

These data suggest that fibroblast populations undergo coordinated functional remodeling involving ECM regulation, oxidative stress, inflammation, and fibroblast-cardiomyocyte communication. Inflammatory-associated PTX3/ICAM1 fibroblasts continued to expand despite declining collagen deposition, and were significantly more prevalent in animals with inducible AF, supporting the concept that this population of fibroblasts is more closely associated with the development of the arrhythmogenic substrate in MR.

Together, these findings provide a molecular framework for understanding atrial remodeling during MR progression and identify distinct fibroblast populations and signaling pathways as potential biomarkers for the arrhythmogenic substrate and therapeutic targets for preventing remodeling before the development of AF.

## Novelty & Significance

### What is known?

- MR promotes left atrial remodeling and increases susceptibility to AF.
- Cardiac fibroblasts are active regulators of extracellular matrix remodeling, inflammation, and electrical signaling in cardiovascular disease.
- The molecular mechanisms driving MR progression and AF substrate formation remain poorly understood.

### What new information does this article contribute?

- This study provides the first single-nucleus transcriptomic characterization of major cardiac cell populations and fibroblast heterogeneity in canine MR.
- Distinct fibroblast populations underwent dynamic remodeling during MR progression.
- Fibroblast population remodeling was associated with AF inducibility.
- This study identified candidate fibroblast-cardiomyocyte signaling pathways.

AF is commonly seen in patients with MR, yet the molecular mechanisms linking structural remodeling to arrhythmia susceptibility remain poorly understood. Using snRNA-seq and RNAscope validation, we identified four transcriptionally distinct fibroblast populations that underwent dynamic remodeling throughout MR progression. Although MR caused progressive left atrial dilation, neither chamber enlargement nor total collagen burden explained AF inducibility. Instead, fibroblast population remodeling and altered fibroblast-cardiomyocyte communication were associated with arrhythmia susceptibility. These findings provide the first molecular atlas of temporal fibroblast remodeling in a canine MR model and identify fibroblast state transitions and intercellular signaling pathways as potential therapeutic targets for preventing arrhythmogenic remodeling.

## Limitations

This study has several limitations. First, the snRNA-seq analysis was performed on a limited number of animals, which may reduce the ability to detect less abundant cell populations and limited statistical power. Although key findings were validated in an independent cohort of 18 canines, larger studies are needed to confirm the reproducibility of these fibroblast populations and their remodeling. Second, this work was performed in a canine MR model, and the extent to which these fibroblast populations and molecular programs are conserved in human MR and AF remains to be determined. Finally, this study identified transcriptionally distinct fibroblast populations and their associations with disease progression and AF inducibility but did not establish their causal roles in arrhythmogenesis. Future studies will be needed to determine how specific fibroblast populations contribute to atrial remodeling and the development of AF.

## Acknowledgments

We thank the Alvin J. Siteman Cancer Center, supported in part by NCI Cancer Center Support Grant P30 CA091842, at Washington University School of Medicine and Barnes-Jewish Hospital in St. Louis, MO, for the use of the Tissue Procurement Core and Genome Technology Access Center, which provided nuclei isolation and single-nucleus RNA sequencing services. We also thank the Institute of Clinical and Translational Sciences (ICTS), funded by the National Institutes of Health’s NCATS Clinical and Translational Science Award (CTSA) Program (UL1 TR002345) for supporting the Genome Technology Access Center. Confocal imaging, purchased with support from NIMH grant MH126964, was performed in part through the Washington University Center for Cellular Imaging (WUCCI), supported by Washington University School of Medicine, The Children’s Discovery Institute of Washington University and St. Louis Children’s Hospital (CDI-CORE-2015-505 and CDI-CORE-2019-813), and the Foundation for Barnes-Jewish Hospital (3770 and 4642). We thank Dr. Michael Vasek and the Dougherty Lab for their RNAscope expertise and use of equipment.

## Sources of Funding

NIH (R01-HL032257) Foundation for Barnes-Jewish Hospital

## Disclosures

S.R. has received research funding for her institution from Varian Medical Systems.

## Supplemental Material

Supplemental Methods Figures S1-S6

Tables S1-S3

Major Resources Table

## Nonstandard Abbreviations and Acronyms

ADRB2: Adrenoceptor Beta 2
AF: Atrial Fibrillation
ECM: Extracellular Matrix
GRIA4: Glutamate Ionotropic Receptor AMPA Type Subunit 4
HCN1: Hyperpolarization Activated Cyclic Nucleotide Gated Potassium Channel 1
ICAM1: Intercellular Adhesion Molecule 1
LA: Left Atrium
LAPW: Left Atrial Posterior Wall
MR: Mitral Regurgitation
NOX4: NADPH Oxidase 4
PCOLCE2: Procollagen C-Endopeptidase Enhancer 2
PTX3: Pentraxin 3
RNAscope: RNA In Situ Hybridization Assay
snRNA-seq: Single-Nucleus RNA Sequencing

